# Reduction of low-frequency oscillations in cerebral circulation correlates with pupil dilation during cognition: an fNIRS study

**DOI:** 10.1101/2023.11.30.569480

**Authors:** Mayra Moreno-Castillo, Elias Manjarrez

## Abstract

Numerous studies have utilized functional near-infrared spectroscopy (fNIRS) to investigate brain hemodynamics during diverse cognitive tasks. However, although these studies have consistently reported increased oxygenation levels in the prefrontal cortex, they have not explored their effects on the amplitude of brain low-frequency oscillations (LFO). Additionally, other reports have shown that pupil dilation occurs during cognitive tasks, which may indicate a correlation between LFO amplitude and pupil dilation. However, no study has demonstrated such a correlation. Our research has two aims: firstly, to analyze the impact of cognitive tasks on the amplitude of LFO using oxyhemoglobin (HbO2) fNIRS signals, and secondly, to assess the relationship between the amplitude of these LFO and pupil diameters during such cognitive tasks. We found that during arithmetic tasks, the brain LFOs recorded on the prefrontal cortex were temporarily reduced while pupil diameter increased. These findings offer new insights into the physiological functions of reactivity of LFO in cerebral circulation. Additionally, combining fNIRS signals to track LFO on the prefrontal cortex with pupil measurement shows the feasibility of developing efficient hybrid brain-computer interfaces of LFO-pupil detection capable of predicting the initiation and ending of cognitive processes.

**Significant statement:** Inspired by the Berger effect, which demonstrates alpha wave reactivity with eye-opening, our study investigated reactivity in other physiological oscillations related to brain hemodynamics during cognition. Our findings revealed a novel phenomenon: a temporary reduction in the amplitude of low-frequency oscillations (LFOs) of oxyhemoglobin (HbO2) in the brain’s prefrontal cortex, concurrent with increased pupil diameter during arithmetic tasks. This discovery expands our knowledge of LFO in cerebral circulation and creates new avenues for studying fNIRS-LFO reactivity during cognitive tasks in healthy individuals and patients.

## Introduction

The cerebral vascular system contains an intricate network of conduits transporting oxyhemoglobin (HbO2) and deoxyhemoglobin (Hb), which are fundamental to the neurophysiological integrity of the brain. This system bifurcates into two critical pathways: 1) arteries, which transport oxygenated blood from the lungs’ alveoli to the neuronal synapses, and 2) veins, which clear metabolic detritus from neural processes. The symbiotic operation of the respiratory and cardiovascular systems supports neuronal functionality, engaging in a sophisticated exchange of signals that includes oxygen supply dynamics, neurovascular coupling, local vasomotion, and the regulatory control of the autonomic nervous system. In this autonomic interface, a systemic modulation of cardiac pulsation and vascular pressure occurs, giving rise to physiological oscillations, notably the cardiac rhythm (∼1 Hz) and the less-frequent Mayer waves (∼0.1 Hz in humans). The Mayer waves are sympathetically mediated LFOs (Julien, 2006, 2020), but other non-sympathetically produced LFOs occur locally in the brain and are generated by vasomotion. These vasomotion LFOs can be reduced in amplitude by an augmentation in local neuronal activity (Brown et al., 2002). In the brain, the LFO Mayer-waves and the vasomotion LFOs, among other unknown oscillatory mechanisms, are exhibited at the frequency range from 0.06 to 0.125 Hz. Therefore, the rationale of our study is to examine changes in the amplitude of LFOs in this frequency range during cognitive tasks. In general, the frequency ranges of the hemodynamic oscillations recorded in the brain with fNIRS are 0.6-2 Hz for cardiac activity, 0.15-0.6 Hz for respiration, 0.06-0.12 Hz for LFO, and 0.01-0.05 Hz for very low-frequency oscillations (VLFO) (Obrig et al., 2000; Taga & Watanabe, 2023). Consistently, fMRI also detects LFOs in blood-oxygen-dependent (BOLD) signals between 0.06-0.125 Hz (Elwell et al., 1999; Rayshubskiy et al., 2013; Murphy et al., 2013; Whittaker et al., 2019).

Continuing with the idea that Mayer waves are linked to sympathetic activity, there is another sympathetically mediated response that has been extensively researched: pupil dilation. Many studies have documented that when the brain is engaged in cognitively demanding tasks, pupils tend to dilate (Haab, 1903; Weiler, 1910; Hess & Polt, 1964; Kahneman & Beatty, 1966; Beatty, 1982; Landgraf et al., 2010; Wierda et al., 2012; Smallwood et al., 2011; Kang et al., 2014; Lisi et al., 2015; Dix & Van der Meer, 2015; Foroughi et al., 2017; O’Neill et al., 2000; Siegle et al., 2023; Kolnes et al., 2023; Mihelčič & Podlesek, 2023; for review see: Viglione et al., 2023; Zenon, 2019; Layzer Yavin et al., 2022).

However, during cognitive tasks, pupil dilation not only occurs, but direct current (DC) HbO2 levels detected with fNIRS also increase, closely linking them to cognitive and neuronal activity (Jobsis, 1977; Ferrari & Quaresima, 2012); for recent reviews (see Zhou et al., 2023; Liu et al., 2023; Hoshi & Tamura, 1993; Menon et al., 2002; Hugdahl et al., 2004; Tanida et al., 2004; Tanida et al., 2007; Yang et al., 2009; Richter et al., 2009; Pfurtscheller et al., 2010; Power et al., 2010; Verner et al., 2013).

Based on the information provided in the previous paragraphs, we can summarize that both the DC levels of HbO2 detected with fNIRS and the size of the pupil increase during cognitive tasks. However, given that both pupil size (Steinhauer et al., 2004; Ferencová et al., 2021; Ishikawa, 2023) and Mayer waves (Julien, 2006, 2020) are controlled by a balance of sympathetic and parasympathetic components, which are responsible for our body’s ‘fight or flight’ responses, it is reasonable to think that these Mayer waves and the vasomotion LFOs mediated by neuronal activity in the cerebral circulation might also change when we are thinking hard. Our study is built around a straightforward hypothesis: while we are engaged in a challenging arithmetic task, as our pupils get larger, the amplitude of LFOs at the frequency range of 0.06 to 0.125 Hz might get smaller. Using fNIRS to track HbO2 levels in the brain, we have set out to test this hypothesis and to see how these changes in LFO amplitude line up with changes in pupil size during a difficult cognitive task. We believe that by looking into how LFOs behave during thought processes, we can contribute to uncovering how the brain regulates its LFO rhythms of blood flow when we are concentrating.

## Materials and Methods

### Participants

The study was conducted on 49 healthy individuals with no history of neurological disease. The study was divided into two protocols. The first protocol involved using fNIRS to track HbO2 levels on the bilateral prefrontal cortex with eyes closed. The second protocol involved using fNIRS to track HbO2 levels on the bilateral prefrontal cortex while monitoring changes in pupil diameter. The recordings were taken before (initial control), during the performance of mathematical tests (Math), and after (final control).

In the first protocol, 26 people participated, but only 20 were included in the analysis. Six were excluded (two experienced drowsiness and fatigue and fell asleep during the mathematical questioning period, while the remaining four had many movement artifacts in their records that made analysis difficult). In the second protocol, 23 healthy individuals took part. Of these, three were excluded (two due to drowsiness and one due to movement artifacts). Additionally, one participant’s pupil images were unclear, making it impossible to measure pupil diameter. All participants agreed to the Declaration of Helsinki and signed an informed consent. A local ethics committee of the Benemérita Universidad Autónoma de Puebla, Mexico, approved the experimental protocol.

### Positioning of the HbO2 fNIRS system on the prefrontal cortex region

We used a custom-made fNIRS system to measure changes in relative oxyhemoglobin (HbO2) concentration in real time. Our method employed the continuous wave (CW) technique and the modified Lambert-Beer’s law with the required extinction coefficient, following the guidelines (do-it-yourself) described by Tsow et al., 2021. We measured in real time the HbO2 fNIRS signal in the bilateral dorsomedial/dorsolateral prefrontal cortex, which is Brodmann area 9 (Croxson et al., 2005). We used a 1/1 arrangement with the photodiode OPT-101 and near-infrared LED, keeping 2.5-3.0 cm between them (Chen et al., 2020). The arrays were placed bilaterally, following the coordinates of the 10-20 system. For the left hemisphere, we selected the quadrant formed by Nasion-FP1-F3-FZ, and for the right hemisphere, the quadrant formed by Nasion-FZ-F4-FP2. These quadrants correspond to the bilateral dorsomedial/dorsolateral prefrontal cortex (Croxson et al., 2005). To ensure the safety of the participants, we monitored the finger partial oxygen saturation (SpO2) and heart rate using a pulse-oximeter device (NONIN 7500). None of the participants experienced any discomfort or showed any alteration in their vital signs during the protocol.

### Experimental protocol of HbO2 fNIRS recording during a cognitive task

The experimental protocol consisted of four blocks, each lasting 120 seconds. Block 1 was the initial control, where the participant remained at rest, sitting in a comfortable position. Block 2, called Math1, presented successive arithmetic operations consisting of two-digit additions of low difficulty. Block 3, called Math2, presented arithmetic operations consisting of four-digit additions of high difficulty. Block 4 was the final control, where participants returned to a resting state. During all blocks, participants kept their eyes closed.

During the cognitive tasks of Math1 and Math2, participants were instructed to mentally solve the arithmetic operations announced by the experimenter. To avoid potential artifacts generated by verbalizing the response, participants were instructed to press a button held in their hands once they had solved the arithmetic question. After pressing the button, the experimenter asked the following questions for four minutes.

The experimental protocol described above was repeated five times (n=5 trials) for each participant.

### Experimental protocol of simultaneous fNIRS recording and pupil size during a cognitive task

In a different set of experiments, we collected both fNIRS recordings and a sequence of pictures of participants’ pupils using a camera without flash to measure their pupil diameter. The protocol was divided into two stages, each lasting two minutes. During the first stage, we obtained the fNIRS recordings and pupil pictures while they sat comfortably. In the second stage, the participants were asked to perform arithmetic sums that started with easy operations and gradually increased in difficulty. During this arithmetic task, we also obtained the fNIRS recordings and pupil pictures. They were instructed to listen carefully to the operations, mentally calculate the answer, and respond by pressing a button. During the experiment, participants underwent two lighting conditions. In one condition, they were exposed to “ambient room light + artificial light turned ON” from a panel placed 40 cm lateral to them. In the other condition, they were only exposed to ambient room light with the artificial light turned OFF. We measured the light intensity on the subjects under both conditions with a Lux meter Hioki FT3424 (Hioki E.E. Corporation, Japan). The light intensity on the subject composed by “ambient light + light panel turned OFF” was 489 ± 53 Luxes. On the other hand, the light intensity on the subject composed by “ambient light + light panel turned ON” was 886 ± 38 Luxes. There was a statistically significant difference (p<0.001) between both light intensities.

### Statistical Analysis

We studied the percentage change of LFO during the Math1 and Math2 cognitive tasks and the initial control condition for 20 participants. Since the LFO data obtained from these subjects were not normally distributed, we used the Kruskal-Wallis one-way analysis of variance on ranks to compare the dependent variable “percentage change of LFO” across the two Math1 and Math2 cognitive tasks. We considered all effects in the Kruskal-Walli’s test significant if p<0.001. Then, we used Dunn’s test for post-hoc testing for multiple comparisons versus the control group. We reported all effects in the Dunn’s test as significant if p< 0.05.

We also compared the percentage change of LFO or the pupil diameter during the Math2 cognitive task and the initial control condition for 19 participants. In this case, the LFO data obtained from these subjects were normally distributed, so we used the student’s t-test to compare the two groups. However, the data was not normally distributed when the condition of the light panel turned OFF. Thus, we used the Mann-Whitney rank sum test to compare the two groups.

To examine whether there is a linear correlation between the percentage change of LFO and the difference in pupil size during the Math2 task compared to the control condition, we used the Pearson product-moment correlation coefficient. The correlation coefficients were calculated for n-2 degrees of freedom (DF). We considered the correlation coefficients significant if p<0.05. Statistical analyses were conducted using SigmaPlot (Systat Software Inc., version 11.0). We also used MATLAB version R2022b developed by MathWorks Inc. in Natick, Massachusetts.

## Results

We conducted a detailed examination of the real-time traces of HbO2 fNIRS signals recorded from the prefrontal cortex on both the right and left sides using a single near-infrared LED. In Figure 1, we present continuous raw examples of two real-time recordings, without averaging or filtering, obtained from a participant. The four stages of the cognitive task are separated by vertical dashed lines, with the first stage representing 120 seconds of the initial control, the second and third stages denoting the application of the mathematical tasks math1 and math2, respectively, for 120 seconds each, and the fourth stage of about 120 seconds corresponding to the final control. The lower panels of Figures 1A and B reveal that during control conditions, slow oscillations between 0.06 to 0.125 are frequently observed, but their amplitude is significantly reduced during the Math1 and Math2 conditions. These waves are within the LFOs range. Overall, we can qualitatively observe that the amplitude of these LFOs is reduced during the Math1 and Math2 conditions.

**Figure 1.**
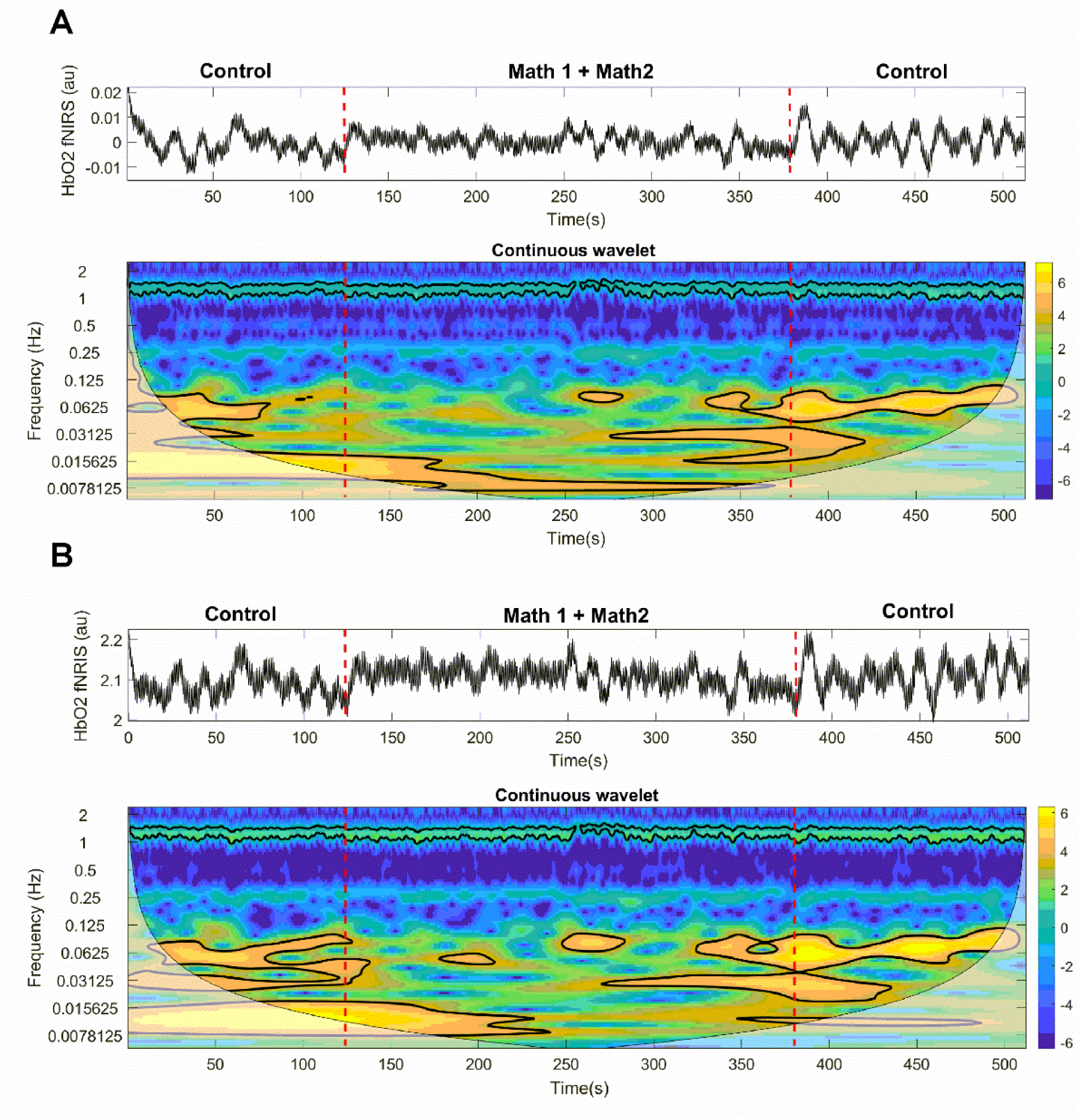
The figure shows continuous raw recordings of HbO2 fNIRS from a participant. The recordings are not averaged or filtered. Recording **A** is obtained from the right prefrontal cortex while recording **B** is obtained from the left prefrontal cortex. The figure is segmented into four stages by vertical dashed lines, which are initial control (120 seconds), Math1 task (120 seconds), Math2 task (120 seconds), and final control (approximately 120 seconds). During control conditions, the lower panels highlight in yellow the presence of slow oscillations (0.06 to 0.125 Hz) within the LFO (low-frequency oscillations) range. These oscillations have a notable reduction in LFO amplitude during the Math1 and Math2 tasks. The oscillations of small amplitude are represented in blue color, while those oscillations of high amplitude are illustrated in yellow color, according to the color scale on the right, which is in arbitrary units. The purpose of this figure is to show the decrease in LFO amplitude during Math1 and Math2 cognitive tasks compared to control conditions.

During our study, we aimed to measure changes in the amplitude of the low-frequency oscillations (LFOs) by analyzing the power spectra of HbO2 fNIRS signals. An example of the power spectral analysis of the HbO2 fNIRS for an individual is shown in Figure 2. We measured the power spectrum amplitude in the LFO range (0.05 to 0.13 Hz) for the right and left prefrontal cortex, respectively, during the four stages of the cognitive task: Initial control, Math1, Math 2, and Final control. The asterisks in the zoom of Figures 2C and D indicate the highest peaks in the power spectra of HbO2 fNIRS signals. The maximal peak frequency of LFO was the frequency at which the highest peak in the power spectrum of HbO2 fNIRS oscillations from 0.05 to 0.13 Hz was observed. Such maximal peak frequency is the dominant frequency of LFO. Notably, the maximal power spectrum amplitude in the LFO range (0.08 to 0.1 Hz) was significantly reduced during the Math1 and Math2 stages compared to the controls. Similarly, we measured the power spectrum amplitude in the LFO range (0.05 to 0.13 Hz) of n=328 trials and n=298 trials for the right and left prefrontal cortex, respectively, performed by all the participants (N=20) during the four stages of the cognitive task: Initial control, Math1, Math 2, and Final control. The results of such analysis are illustrated in the pooled data in Figure 3. Data are expressed as percentage changes of LFO amplitude relative to the initial control versus the dominant frequency of LFO. The Kruskal-Wallis one-way ANOVA of ranks of data from Figure 3A showed a significant reduction (H=115.9 with 3 degrees of freedom, p<0.001, n=328, Cohen’s d=1.4) in the LFO amplitude during the Math1 and Math2 stages of the cognitive task compared to the initial control stage. Moreover, we performed post-hoc testing for multiple comparisons versus the control group with Dunn’s test, and we found that the multiple comparisons were statistically different (p<0.05). Similarly, the Kruskal-Wallis one-way ANOVA of ranks of data from Figure 3B showed a significant reduction (H=75.0 with 3 degrees of freedom, p<0.001, n=298, Cohen’s d=1.1) in the LFO amplitude during the Math1 and Math2 stages of the cognitive task compared to the initial control stage. Moreover, we performed post-hoc testing for multiple comparisons versus the control group with Dunn’s test, and we found that the multiple comparisons were statistically different (p<0.05).

**Figure 2.**
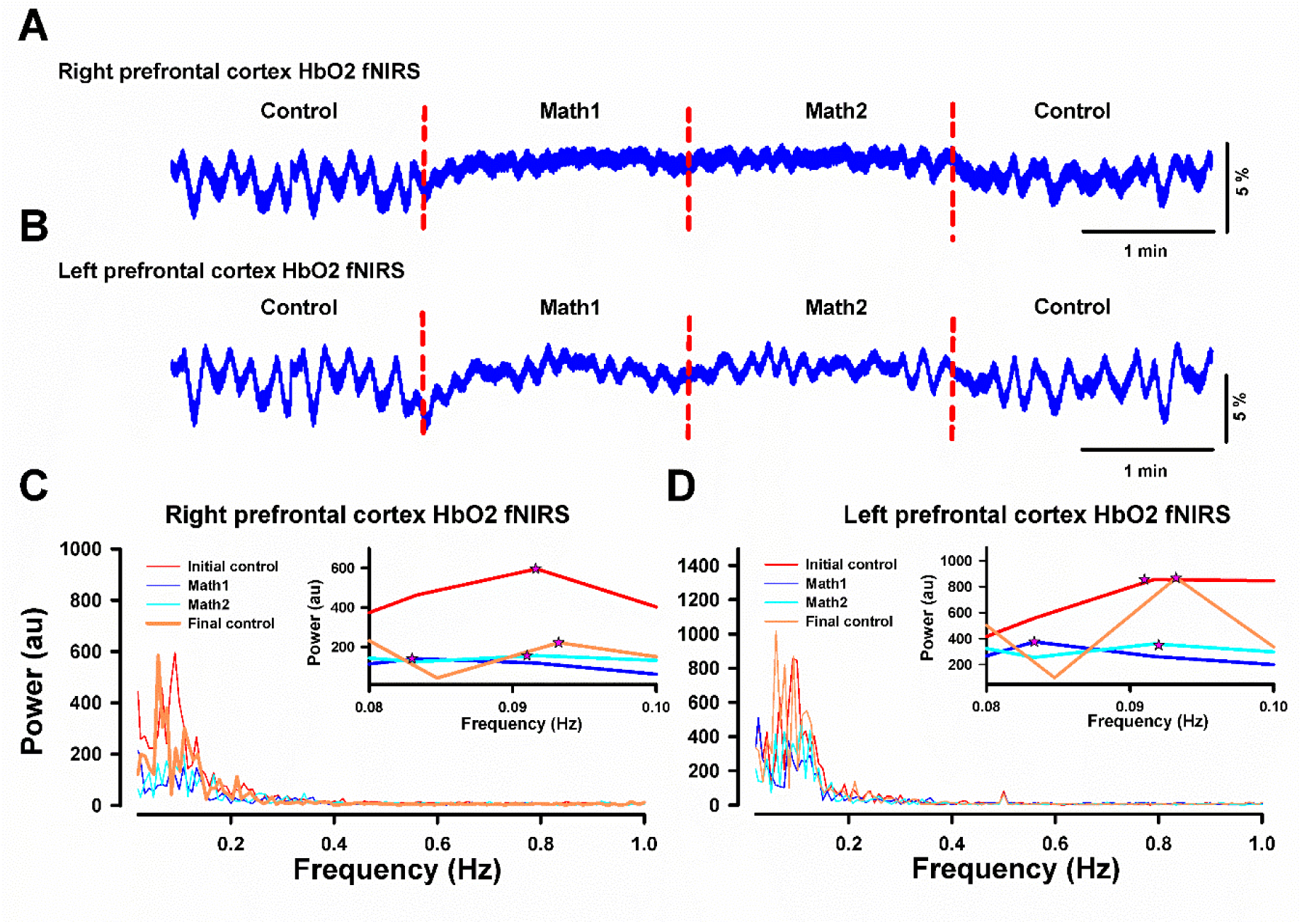
The aim of this figure is to explain how to measure the amplitude of LFO (low-frequency oscillations) from the power spectra of HbO2 fNIRS signals. Data from the right (**A**) and left (**B**) prefrontal cortex obtained from a single participant is used to illustrate the procedure. The power spectra from signals in A and B are shown in graphs **C and D,** respectively. The insets in **C and D** highlight the highest peaks in the power spectrum of HbO2 fNIRS oscillations, marked with asterisks. The highest peaks of these LFOs occur in dominant frequencies (maximal peak frequencies) within the range from 0.08 to 0.1 Hz. The analysis shows that the LFOs in the range from 0.08 to 0.1 Hz are significantly reduced during Math1 (blue color) and Math2 (green color) stages compared to control conditions (red and orange colors, respectively). This procedure was used to create the graphs in Figure 3.

**Figure 3.**
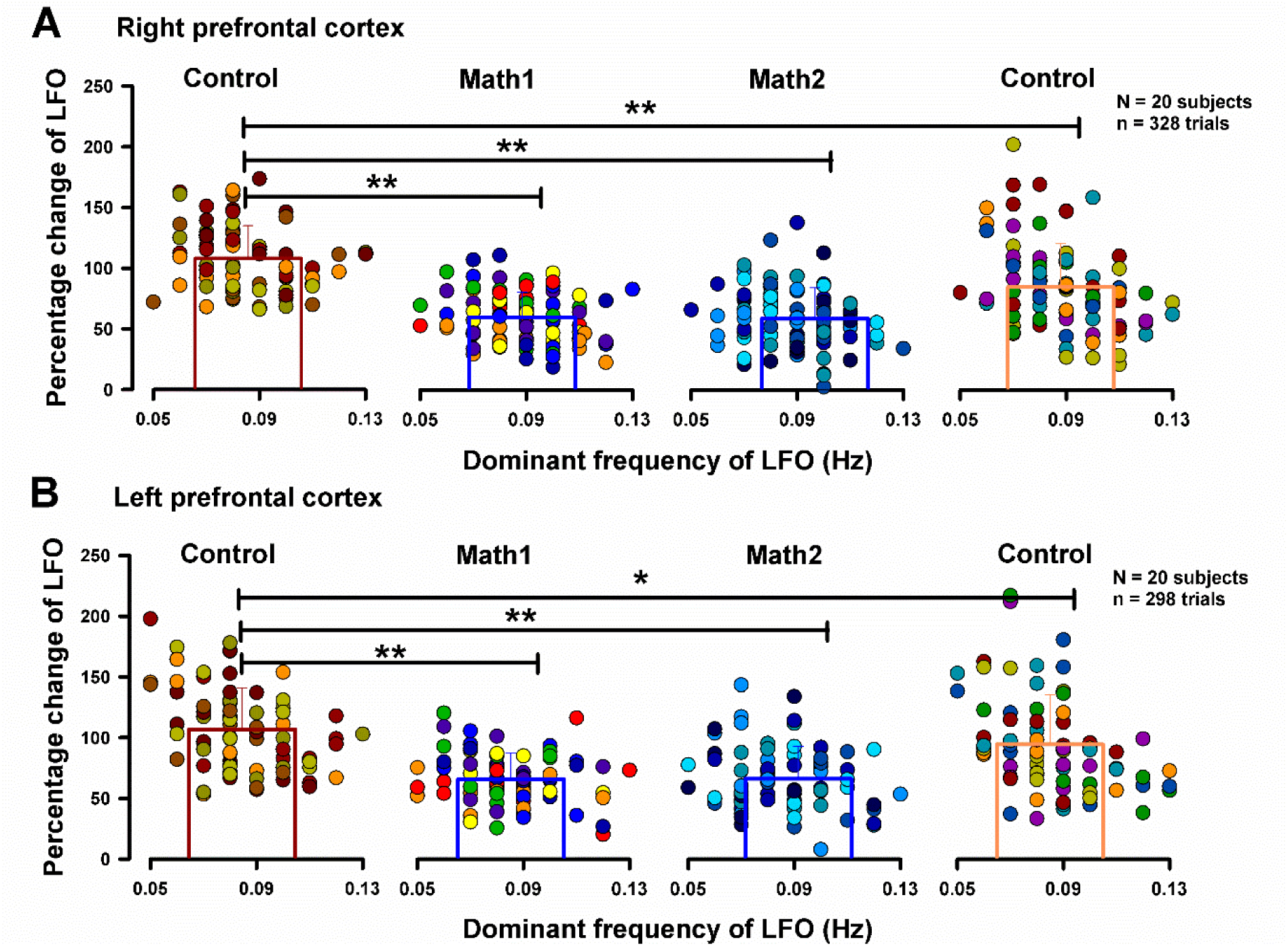
Pooled data illustrating the percentage change of LFO (low-frequency oscillations) relative to the initial control, focusing on the dominant frequency of LFO, as illustrated in Figure 2. **A**, for the right prefrontal cortex. **B**, for the left prefrontal cortex. Data was collected from N=20 participants during the four stages of the cognitive task: Initial control, Math1, Math2, and Final control. Statistical analysis reveals a significant reduction (***p<0.001) in the LFO during the Math1 and Math2 stages compared to the control stages. Please refer to the text to obtain more information about the statistical comparisons.

In another series of experiments, we simultaneously measured changes in pupil diameter and LFO amplitude during the control stage and the Math2 stage. Figure 4 provides an example of raw data obtained from one participant, depicting the behavior of their LFOs and pupil diameters during control (Figure 4A) and Math2 (Figure 4B) stages. These figures qualitatively show that the amplitude of LFOs reduces during the Math2 task while the pupil diameter increases.

**Figure 4.**
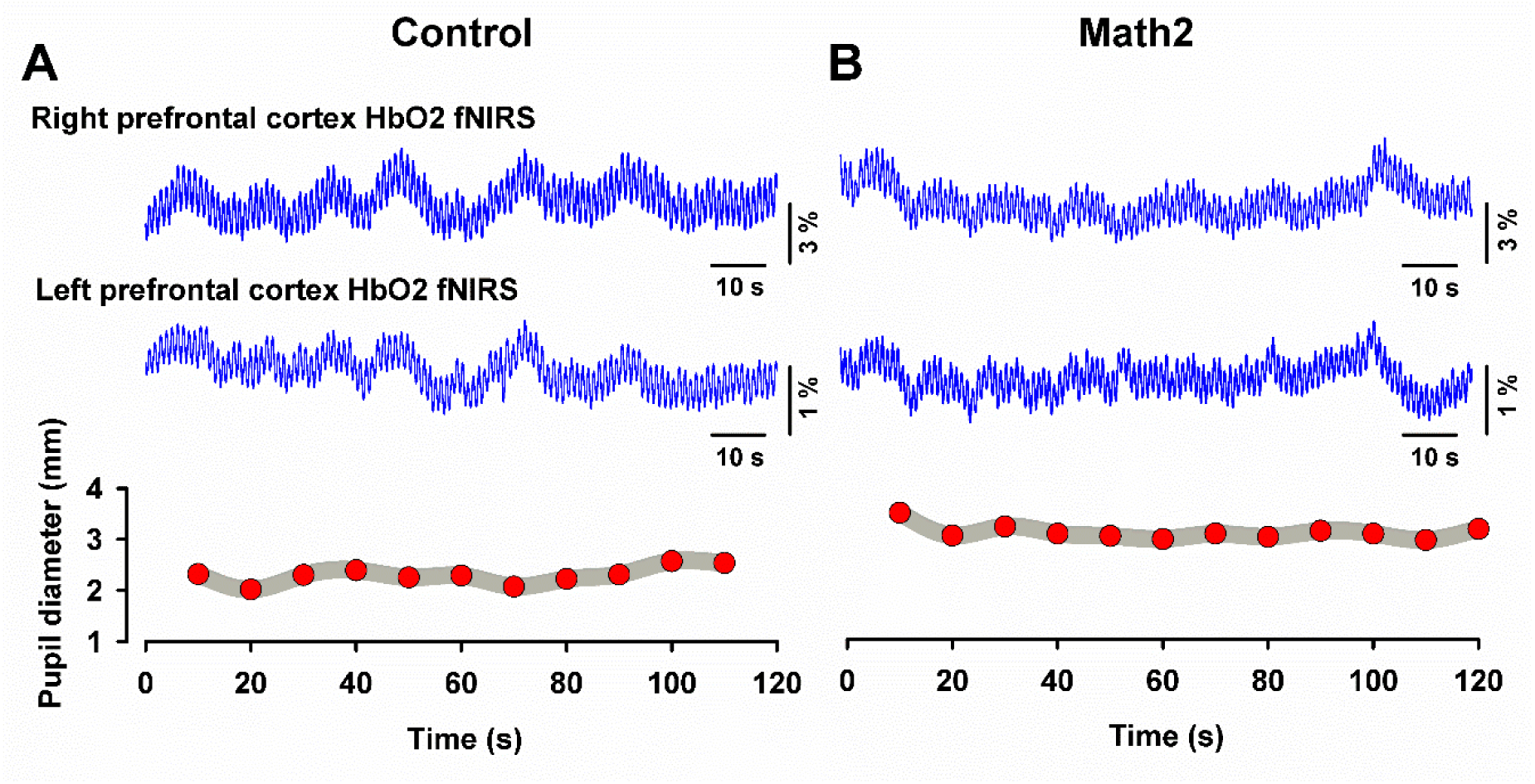
The purpose of this figure is to show that during the Math2 cognitive task (panel B), the LFO (low-frequency oscillations) amplitude decreased while pupil diameter increased compared to the control condition (panel **A**), using data from a single participant.

We analyzed in more detail whether the increment of pupil diameter during the cognitive task Math2 is modulated by the light intensity during the task in 19 participants. We measured the pupil diameter in 12 consecutive pupil pictures during the control condition and another 12 consecutive pupil pictures during the Math2 cognitive task, as illustrated in Figure 5. Figure 5 shows the measured pupil diameters in 19 participants in two lighting conditions: 1) with room light + a light panel turned ON (Figure 5A) and with room light and a light panel turned OFF (Figure 5B). This figure illustrates that the augmentation of pupil diameter during the Math2 task was possible regardless of whether the light panel was turned ON or OFF. We compared the pupil diameter from the control and the Math2 conditions in Figure 5A. Because the data was not normally distributed, we performed a Mann-Whitney rank sum test (n=213 in the control group, n=213 in the Math2 group, U=13793); we found a statistically significant increase in the pupil diameter (p<0.001; Cohen’s d=0.72). Similarly, we compared the pupil diameter from the control and the Math2 conditions in Figure 5B. Because the data was not normally distributed, we performed a Mann-Whitney rank sum test (n=226 in the control group, n=226 in the Math2 group, U=39954); we also found a statistically significant increase in the pupil diameter (p<0.001; Cohen’s d=1.1).

**Figure 5.**
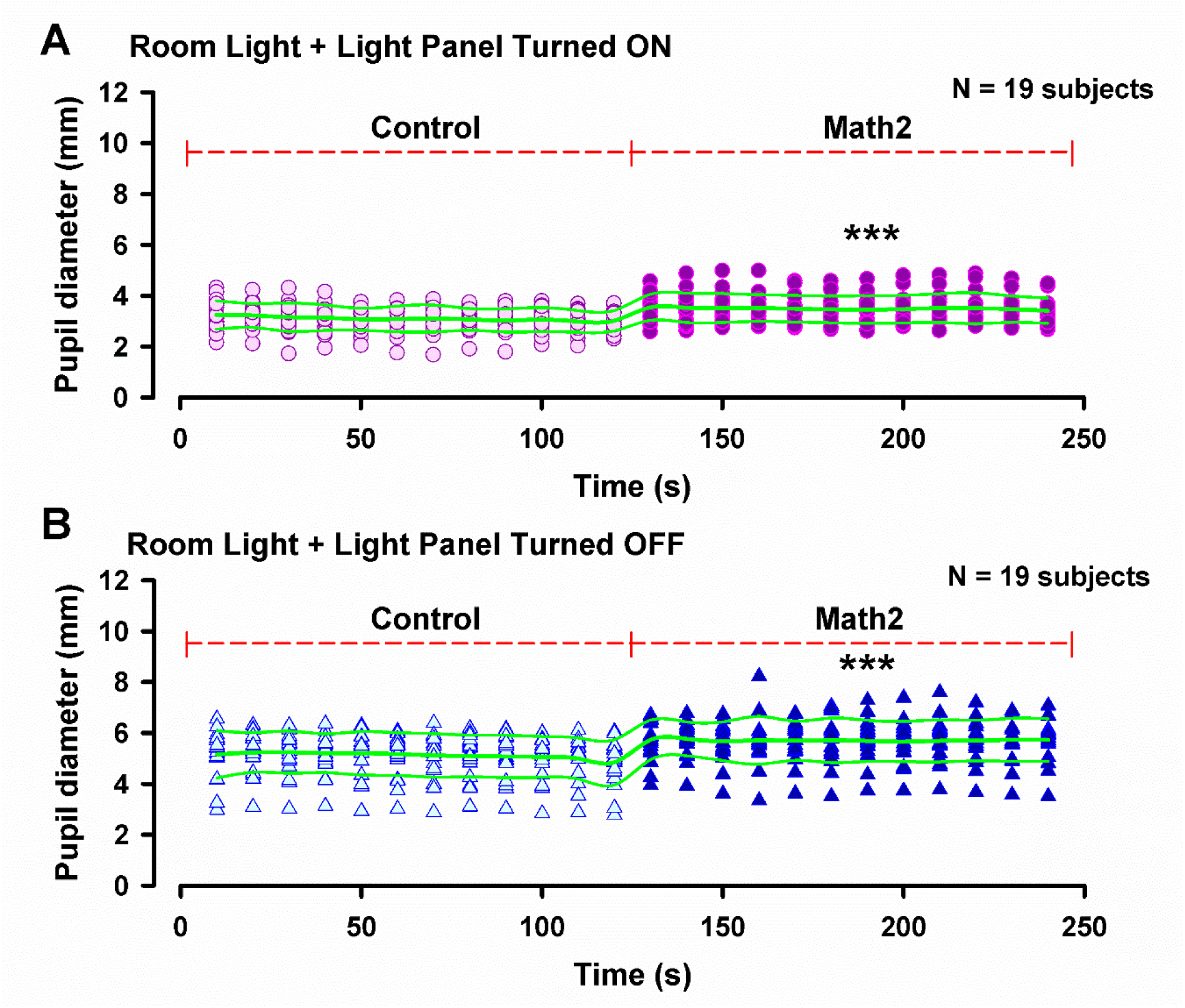
The dataset includes the results obtained from 19 individuals, showing the changes in pupil diameter during the control and the Math2 cognitive task, in different light conditions. Graph **A** shows the data with the light panel turned ON, along with the room light, whereas Graph **B** represents the data with the room light and the light panel turned OFF. These graphs demonstrate an increase in pupil diameter during the Math2 task (*** p<0.001), irrespective of whether the light panel was turned ON or OFF under room light. The graphs also show that in the “light panel turned ON” condition, the pupil diameters exhibit reduced absolute values (during Control and Math2) relative to the “light panel turned OFF” condition.

Our findings indicate that the LFO amplitude in 19 individuals during the cognitive Math2 task was significantly reduced for both illumination conditions (as depicted in Figures 6A, 6B, and 6D, 6E). The student’s t-test data from Figure 6A showed a significant decrease (t=7.1 with 60 degrees of freedom, p<0.001, n=32, mean=101.3, SD=21.8 for the control group and n=30, mean=65.1, SD=17.9 for the Math2 group, Cohen’s d=1.8) in the LFO amplitude during the Math2 stage of the cognitive task compared to the initial control stage. Furthermore, a significant decrease in LFO amplitude was also observed for the left prefrontal cortex (as illustrated in Figure 6B). The student’s t-test data from Figure 6D showed a significant reduction (t=7.0 with 55 degrees of freedom, p<0.001, n=31, mean=100, SD=19 for the control group and n=26, mean=64.5, SD=18.5 for the Math2 group, Cohen’s d=1.8) in the LFO amplitude during the Math2 stage of the cognitive task compared to the initial control stage. Similarly, a significant decrease in LFO amplitude was observed for the left prefrontal cortex (as shown in Figure 6E).

**Figure 6.**
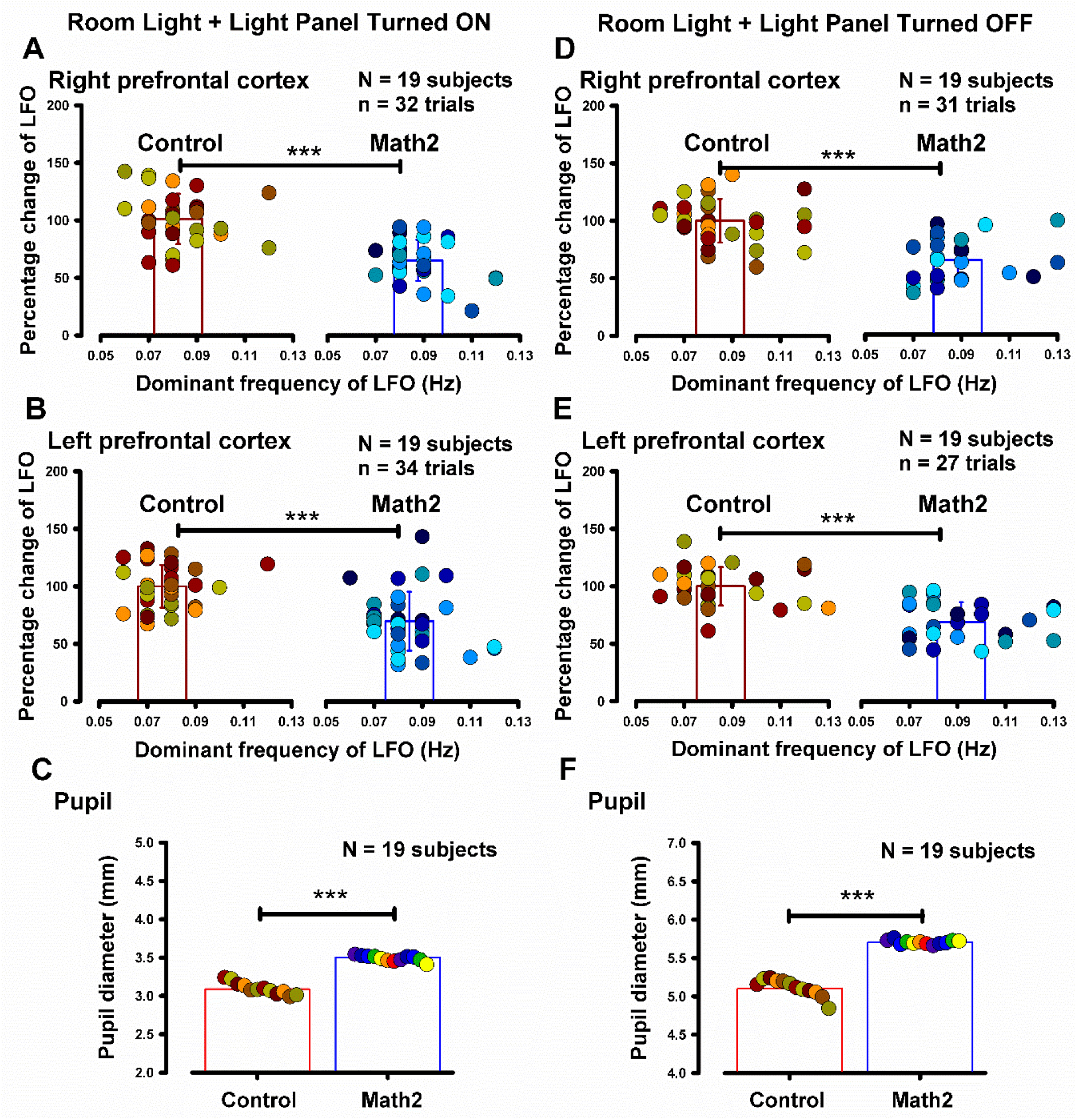
Pooled data of 19 participants showing the percentage change of LFO (low-frequency oscillations) and pupil diameter during the Control and the Math2 cognitive task under the two illumination conditions shown in Figure 5. Condition in **A** represents the room light plus the light panel turned ON, while Condition in **B** represents the room light plus the light panel turned OFF. All the figures indicate a statistically significant reduction (***p<0.001) in the LFO amplitude during the Math2 task compared to the Control under both lighting conditions. In addition, figures **C** and **F** demonstrate an increase in pupil diameter during the Math2 task in both illumination conditions compared to the control.

Additionally, we found that in both illumination conditions, the means of the pupil diameter for the 19 individuals increased during the Math2 condition compared to the control (Figures 6C and 6F). Figures 6C and 6F represent the mean pupil diameters (obtained from 12 pupil pictures of 19 subjects) from Figures 5A and 5B. The student’s t-test of data from Figure 6C showed a significant increase (t=-15.6 with 22 degrees of freedom, p<0.001, n=12, mean=3.0, SD=0.07 for the control group and n=12, mean=3.4, SD=0.03 for the Math2 group) in the pupil diameter during the Math2 stage of the cognitive task compared to the initial control stage. A similar significant increase in pupil diameter was found for the left prefrontal cortex (Figure 6F).

The results presented in Figures 7A and 7B show a significant negative correlation (r=-0.57/DF=27 and r=-0.58/DF=44, p<0.0011) between the percentage of change of LFO from control to Math2 condition and the difference in pupil diameter when the light panel was turned on, in both the right and left prefrontal cortex. However, there was no statistically significant correlation (p>0.2) when only the room illumination was presented, as shown in Figures 7C and 7D (r=-0.25/DF=33, and r=-0.14/DF=30 for the right and left prefrontal cortex, respectively).

**Figure 7.**
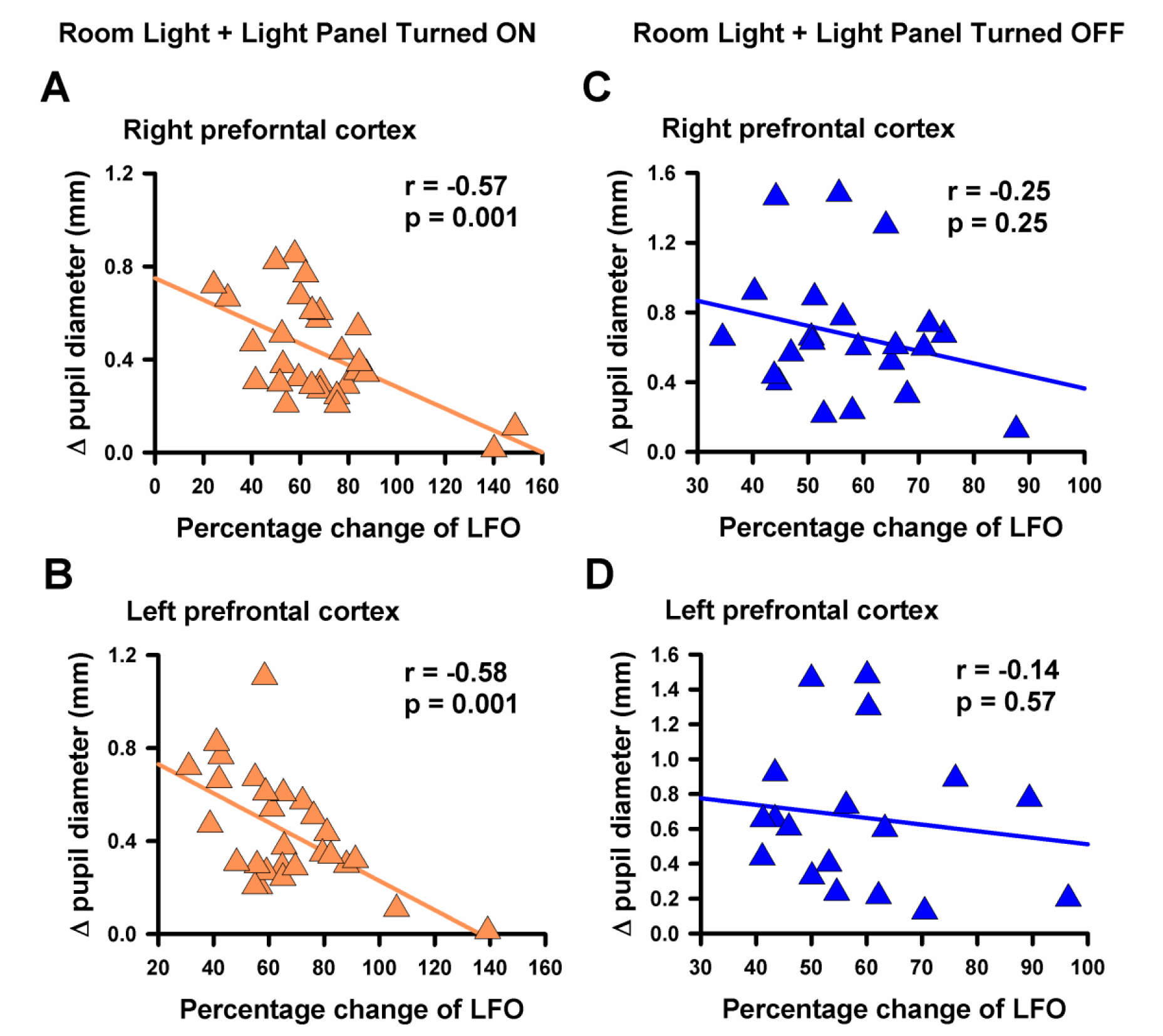
The data shows how the percentage change of LFO (low-frequency oscillations) compared to the initial control correlates with the increase in pupil diameter for the 19 participants analyzed in Figure 6. The results are separated into two sets: **A** and **B**, representing the right and left prefrontal cortex, respectively, during the condition of “room light + light panel turned ON” and **C** and **D**, the same as A and B, but for the condition of “room light + light panel turned OFF”. It’s important to note that adding a light source helps create a statistically significant correlation between the percentage change of LFO relative to the initial control and the increment in pupil diameter. The correlation coefficients were found to be –0.57 and –0.50 for the right and left prefrontal cortex, respectively, with a p value of less than 0.001.

## Discussion

Our research has found that when engaging in cognitive tasks, the amplitude of LFOs in cerebral circulation is reduced in correlation with pupil dilation. Since pupil dilation is controlled by sympathetic activation, these findings provide new insights into the activity patterns of LFOs in cerebral circulation when the sympathetic nervous system is in operation. It is important to note, however, that our study only shows a correlation between the amplitude reduction of LFOs and pupil dilation and not a causal relationship. Therefore, we should exercise caution and not solely attribute the reduction in the amplitude of LFOs during cognitive tasks to the sympathetic nervous system. The value of our findings is that they expand our knowledge of LFO in cerebral circulation and create new avenues for studying fNIRS-LFO reactivity during cognitive tasks in healthy individuals and patients.

It is possible that various mechanisms, in addition to the sympathetic influence, could contribute towards reducing the amplitude of LFO in the brain during cognitive tasks. For example, Morita-Tsuzuki et al., 1992, found that during hypercapnia, the vasomotion LFO in the rat brain decreased in both frequency and amplitude. In contrast, during hypocapnia, their frequency did not change, but their amplitude increased. Similar results were found in humans by Obrig et al. 2000, who reported that hypercapnia attenuates the LFO in fNIRS recordings. This is supported by previous evidence that vasomotion LFOs, which can be generated without autonomic influence, can be attenuated through various physiological factors such as the increase in neuronal activity (Brown et al., 2002) or by electrical stimulation (Filosa et al., 2004). In this context, it is tempting to speculate that the reduction of LFO during cognitive tasks we found here could be due to increased neuronal activity in the prefrontal cortex during the cognitive tasks and to the associated physiological hypercapnia locally occurring in the prefrontal cortex. Hence, we cannot solely attribute the reduction of LFOs during the cognitive task to the Mayer waves. This is because the LFOs in the range of 0.05 to 0.12 Hz may include other oscillations, such as vasomotion-mediated LFOs (Aalkjaer et al., 2011; Sassaroli et al., 2012) in addition to the sympathetically mediated Mayer waves that occur at around 0.1 Hz frequency oscillation.

The ∼0.1 Hz LFOs were first described by Mayer (1876), and according to Julien, the term “Mayer wave” should be reserved for those sympathetically mediated low-frequency oscillations that are slower than respiration (Julien 2006, 2020; see also Seydnejad & Kitney, 2001). Conversely, LFOs produced by local vasomotion oscillations (spontaneous contraction and relaxation of brain blood vessels), occur in the same range of 0.05 to 0.12 Hz but can be produced without sympathetic or parasympathetic influence (Morita et al., 1994; 1995; Sassaroli et al., 2012; Brown et al., 2002; Filosa et al., 2004). Therefore, to investigate the underlying mechanisms of LFOs, it will be necessary to differentiate between the sympathetically mediated Mayer waves and the non-sympathetically mediated vasomotion LFOs in the cerebral circulation. The Mayer waves result from variations in neural discharge in the autonomous nervous system, leading to synchronous Mayer wave oscillations throughout the body. Conversely, vasomotion is a local phenomenon that can persist without neural activity (Nilsson & Aalkjaer, 2003; Aalkjaer et al., 2011; Pradhan & Chakravarthy, 2011; Sassaroli et al., 2012).

On the other hand, it has been shown in animal preparations that when the sympathetic nervous system is activated, the amplitude of Mayer waves may either decrease (Julien et al., 2020) or increase (Julien et al., 2006; Kanbar et al., 2007). This suggests that the amplitude of Mayer waves cannot be solely used to indicate increased vascular sympathetic tone (Julien et al., 2020). Our findings align with these observations, as the cognitive task that led to pupil dilation activated the sympathetic nervous system. This resulted in a decrease in the amplitude of oscillations in the Mayer wave range (∼0.1 Hz), including a decrease in the LFO amplitude. Therefore, we could suggest that the autonomic nervous system may also play a role in reducing the amplitude of the LFO and Mayer waves during the cognitive task.

The concept of modulating the LFOs using physiological processes is not new. Several fNIRS studies have shown increased LFO during motor or sensory interventions. For example, Bajaj et al., 2014 found that during a motor task in humans, the amplitude of LFO increased in HbO2 fNIRS recordings. In another study, Fernandez-Rojas et al., 2019, reported that acupuncture manipulations produced robust LFO activations in the somatosensory cortex. Additionally, Bicciato et al., 2021, found that the amplitude of spectral power in the LFO region of the HbO2 fNIRS recordings increased in all subjects after exposure to familiar music. These studies support the observation that the LFO is susceptible to modulation. However, we found a reduction in the amplitude of the LFO during a cognitive task instead of an amplification. Therefore, comparing our results with those from the mentioned authors, we can conclude that the LFO can be amplified during sensory and motor tasks; conversely, during an arithmetic task like ours, the LFO amplitude is reduced. This comparison suggests that the LFOs represent a physiological observable that could provide insight for developing new research lines into the hemodynamic modulation of the brain during perception, motor action, and cognition.

The physiological interaction between the sympathetic and parasympathetic divisions of the autonomic nervous system controls pupil size. This measurement of pupil size and its responsiveness is known as pupillometry. It is widely used as an indirect marker of brain activity (for review, see van der Wel & van Steenbergen, 2018). Pupillometry is attractive due to its non-invasiveness and the relative ease of recording changes in pupil size. The phenomenon of pupil dilation in response to brain activity was first observed by the German ophthalmologist Haab in 1903. He noticed that the pupils of his patients would dilate when they were asked to perform mental calculations or recall memories. He called this phenomenon the “Hirnrindenreflex” or the “cortical reflex” of the pupil, implying that the cerebral cortex mediated it. Since then, many studies have documented that when the brain is engaged in cognitively demanding tasks, pupils tend to dilate. Here we add new insights to the relationships between pupil dilation and cognition, demonstrating a clear correlation between pupil dilation and the amplitude of the LFO contained in the HbO2 fNIRS recordings. Our results suggest that reducing the pupil diameter with the light panel turned ON can improve the measurement of changes in pupil diameter during a cognitive task and increase its correlation with the LFO amplitude detected in HbO2 fNIRS recordings. A pupil with an already enhanced diameter due to a partially dark room cannot expand further during the cognitive task. Therefore, adding light to reduce the pupil diameter can increase the spatial range in which the pupil diameter can expand during the cognitive task. In this context, our research shows that if we want to examine correlations between pupil dilation and amplitude of LFO, it will be necessary to illuminate the pupil.

We conclude that our findings provide new insights into the reactivity of physiological oscillations in the cerebral circulation, which, like the Berger reactivity, is also a common feature of the human brain. Furthermore, our study demonstrates the feasibility of developing efficient hybrid brain-computer interfaces for LFO-pupil detection by combining fNIRS signals to track LFO on the prefrontal cortex and measuring pupil changes. These interfaces could potentially predict the start and end of cognitive processes.

## Acknowledgments

The following grants supported this research: Cátedra Marcos Moshinsky (E.M) and VIEP-BUAP (E.M).

## Author contributions

Author contributions: E.M. conceived and designed the experiments. M.M.C. and E.M. performed experiments. M.M.C. and E.M. performed the data analysis. E.M. wrote the manuscript. All the authors revised and approved the manuscript.

